# SARS-CoV-2 cell-to-cell spread occurs rapidly and is insensitive to antibody neutralization

**DOI:** 10.1101/2021.06.01.446516

**Authors:** Laurelle Jackson, Hylton Rodel, Shi-Hsia Hwa, Sandile Cele, Yashica Ganga, Houriiyah Tegally, Mallory Bernstein, Jennifer Giandhari, COMMIT-KZN Team, Bernadett I. Gosnell, Khadija Khan, Willem Hanekom, Farina Karim, Tulio de Oliveira, Mahomed-Yunus S. Moosa, Alex Sigal

**Affiliations:** Africa Health Research Institute, Durban, South Africa; Division of Infection and Immunity, University College London, London, UK; School of Laboratory Medicine and Medical Sciences, University of KwaZulu-Natal, Durban, South Africa; KwaZulu-Natal Research Innovation and Sequencing Platform, Durban 4001, South Africa; Department of Infectious Diseases, Nelson R. Mandela School of Clinical Medicine, University of KwaZulu-Natal, Durban 4001, South Africa; Centre for the AIDS Programme of Research in South Africa, Durban 4001, South Africa; Department of Global Health, University of Washington, Seattle, USA; Max Planck Institute for Infection Biology, Berlin, Germany

## Abstract

Viruses increase the efficiency of close-range transmission between cells by manipulating cellular physiology and behavior, and SARS-CoV-2 uses cell fusion as one mechanism for cell-to-cell spread. Here we visualized infection using time-lapse microscopy of a human lung cell line and used live virus neutralization to determine the sensitivity of SARS-CoV-2 cell-to-cell spread to neutralizing antibodies. SARS-CoV-2 infection rapidly led to cell fusion, forming multinucleated cells with clustered nuclei which started to be detected at 6h post-infection. To compare sensitivity of cell-to-cell spread to neutralization, we infected either with cell-free virus or with single infected cells expressing on their surface the SARS-CoV-2 spike protein. We tested two variants of SARS-CoV-2: B.1.117 containing only the D614G substitution, and the escape variant B.1.351. We used the much smaller area of single infected cells relative to infection foci to exclude any input infected cells which did not lead to transmission. The monoclonal antibody and convalescent plasma we tested neutralized cell-free SARS-CoV-2, with the exception of B.1.351 virus, which was poorly neutralized with plasma from non-B.1.351 infections. In contrast, cell-to-cell spread of SARS-CoV-2 showed no sensitivity to monoclonal antibody or convalescent plasma neutralization. These observations suggest that, once cells are infected, SARS-CoV-2 may be more difficult to neutralize in cell types and anatomical compartments permissive for cell-to-cell spread.

## Introduction

If exposure to a virus leads to some cellular infection, infected cells may infect other cells by interacting with them. This happens with multiple virus types [1]. In HIV, cell-to-cell spread involves a virological synapse [2, 3] and is less sensitive to antiretroviral therapy [4] and neutralizing antibodies [5, 6]. A low level of inhibitor was shown to be able to clear infection before but not after the first cells were infected [7].In SARS-CoV-2 infection, the virus causes major changes in cytoskeleton regulation [8] and cell fusion has been observed to be a mechanism of cell-to-cell spread in vitro [9].Fusion is mediated by the binding of the SARS-CoV-2 spike glycoprotein on the cell surface with the cellular angiotensin-converting enzyme 2 (ACE2), the target of spike, on yet uninfected cells. This process is restricted by the interferon pathway and enhanced by the host TMPRSS2 serine protease [9]. Furthermore, spike has two subunits (S1 and S2) with a host furin protease required for cleavage which results in S protein activation. The cleavage of the S1/S2 site has been reported to be necessary for SARS-CoV-2 mediated formation of cellular syncytia and infection of lung cells [10]. Syncytia are one hallmark of SARS-CoV-2 pathology [11, 12, 13], showing that this mechanism of cell-to-cell spread operates in vivo in some environments.

Antibody mediated neutralization of SARS-CoV-2 is a key immune response in Covid-19 disease and is predictive of immune protection [14, 15], with 50% protection from infection achieved at about 20% of mean convalescent antibody levels [14]. The SARS-CoV-2 spike on the virion binds to cell surface ACE2 through the receptor binding domain (RBD) region of spike [16]. The RBD is the main target of neutralizing antibodies elicited by SARS-CoV-2 infection, with additional neutralizing antibodies directed at the N-terminal domain (NTD) of spike [17, 18].

South Africa had two SARS-CoV-2 infection waves to date, with a possible third wave starting as this work is written (https://coronavirus.jhu.edu/map.html). The first wave (peak July 2020) consisted of viral variants with the D614G mutation (referred here as D614G). These variants were replaced by the B.1.351 variant in the second South African infection wave (peak January 2021). This variant has the L18F, K417N, E484K, and N501Y mutations in the NTD and RBD of spike, as well as a deletion in the NTD [19]. B.1.351 has been shown to escape neutralization by convalescent plasma elicited by non-B.1.351 infections, as well as reduce the potency of vaccine elicited neutralization [20, 21, 22, 23, 24, 25, 26, 27, 28, 29, 30]. However, plasma immunity elicited by B.1.351 infection can effectively cross-neutralize the earlier variants which express the D614G mutation but not the cardinal mutations of B.1.351 [20, 31]. The live virus neutralization assay (LVNA) performed by us and by others uses the African green monkey VeroE6 cell line [32]. While this cell line can be used to read out the number of SARS-CoV-2 infectious units, how well it reproduces human cell behavior in SARS-CoV-2 transmission between cells is currently not well understood.

Here, we used a human lung cell line to visualize SARS-CoV-2 cell-to-cell spread by timelapse microscopy and investigate the effect of cell-to-cell spread on the sensitivity of SARS-CoV-2 infection to neutralizing antibodies, where infection was with either with a D614G or the B.1.351 variant.

SARS-CoV-2 was able to rapidly induce cell fusion resulting in multi-nuclear cellular syncytia with clustered nuclei. Cell-free SARS-CoV-2 infection was neutralized with a monoclonal antibody and both D614G-elicited and B.1.351-elicited convalescent plasma, although in agreement with previous results B.1.351 virus was poorly cross-neutralized by D614G-elicited plasma. In contrast, neither the mAb, nor any plasma/variant combination, was effective at reducing cell-to-cell spread of SARS-CoV-2. These results may indicate that SARS-CoV-2 may use cell-to-cell spread to reduce sensitivity to neutralizing antibodies, which may delay viral clearance in environments permissive for this infection mode.

## Results

### Rapid formation of syncytia upon SARS-CoV-2 infection

We engineered a human lung cell line which can be productively infected by SARS-CoV-2 and where cell nuclei can be visualized by fluorescent microscopy. We used the H1299 non-small lung cell carcinoma line, labelled using a CD-tagging approach [33]. We used a clone of the cell line with a yellow fluorescent protein (YFP) labeled histone, leading to fluorescent labelling of the nucleus without overexpression of the fluorescent tag. We then stably expressed the ACE2 receptor in these cells and selected cell clones which were infectable with SARS-CoV-2 (H1299-ACE2, Figure S1).

We performed time-lapse microscopy of live SARS-CoV-2 infection in Biosafety level 3 containment to visualize and quantify cell-to-cell spread of SARS-CoV-2. Addition of a suspension of single infected cells to uninfected cells led to the formation of syncytia (Figure 1A top panel, Movie 1) which did not occur in the absence of infection (movie 2). Syncytia showed distinct nuclear organization, with nuclei often becoming clustered in a ring (Figure 1A, middle and bottom panels. movie 3). This was not observed in uninfected cells even at high cell density (movie 4). We used automated image analysis to detect clustered nuclei (Figure S2). The analysis showed that clustering of nuclei within a multi-nucleated cell occurred in a subset of cells upon SARS-CoV-2 infection, reaching about 20% of nuclei by 36 hours post-infection (Figure 1B). Unlike uninfected cells, there was little cell division in the infected cell culture, as shown by the constant number of nuclei detected over time (Figure 1C). The process of cell syncytia formation was detectable approximately 6 hours post-infection, consistent with the rapid infection cycle of this virus [34].

**Figure 1:**
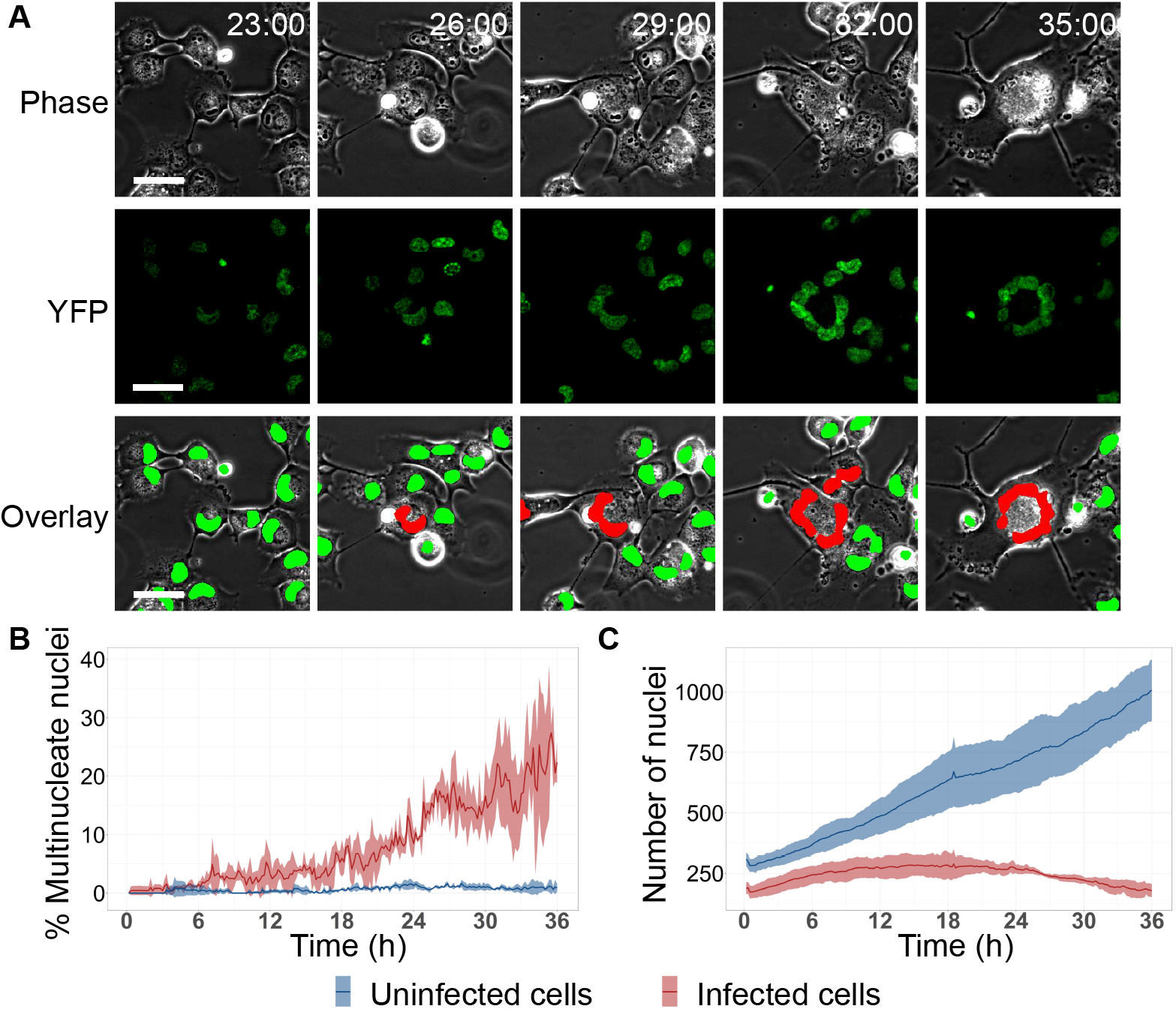
SARS-CoV-2 infected cells fuse to form multinucleated cellular structures with clustered nuclei. (A) Phase contrast (first row), YFP fluorescence (second row), and overlay images (third row) of infected cells, where cell nuclei express YFP. Red in the overlay image denotes nuclei detected as clustered by automated image analysis, while green denotes nuclei detected as non-clustered. Time of image post-infection is on the top right of the phase image as hours:minutes, bar is 50*μ*m. (B) Percent of nuclei in multi-nucleate cells as a function of time. Shown are the mean and standard deviation for infected cells (red) versus uninfected cells (blue). (C) Number of nuclei through time in the timelapse movie. Shown are the mean and standard deviation of the absolute number of nuclei per experiment. Three independent experiments were performed, where each experiment used a different clone of H1299-ACE2 cells.

### SARS-CoV-2 cell-to-cell spread loses sensitivity to neutralization by a monoclonal antibody

To compare SARS-CoV-2 cell-free infection versus cell-to-cell spread, we used as the infection source either filtered supernatant from infected VeroE6 cells, or a single cell suspension of H1299-ACE2 cells infected with the filtered VeroE6 cell supernatant. Virus used was an isolate from the B.1.1.117 lineage which contains the D614G substitution [20] and is referred to here as D614G.

For cell-to-cell spread, cells were infected by cell-free virus for 16 hours before they were used as the infecting donor cells (Materials and Methods). We validated that spike was expressed on the cell surface of input infected cells by anti-spike antibody labelling of non-permeabilized cells (Figure S3), and that we were infecting with single infected cells (Materials and Methods and see below). Allowing the cell suspension to stand for 5 minutes after trypsinization and re-suspension of cells in media was sufficient to separate cell syncytia from the single infected cells to be used as the infection source. We used a live virus neutralization assay (LVNA), quantified by counting the number of focus forming units in a multiwell plate [32, 20] in H1299-ACE2 cells. These cells had well defined foci that were similar to those observed in African Green Monkey VeroE6 cells generally used for LVNA, except that the H1299-ACE2 cells were more sensitive to infection with the same stock of virus (Figure S4). Interestingly, in contrast to VeroE6 cells, focus size did not increase with B.1.351 relative to D614G infection in H1299-ACE2 cells.

We first used the Genscript BS-R2B2 rabbit monoclonal neutralizing antibody (referred to here by its catalog number, A02051) to neutralize infection (Figure 2). We have previously shown that this monoclonal antibody effectively neutralized cell-free virus [20]. We mixed the virus or infected cells with serially diluted antibody for 1 hour, then added the mixture to H1299-ACE2 cells and counted infection foci after 24 hours using automated image analysis (Figure 2A, Materials and Methods). We visualized the infecting cells 2 hours post-addition (Figure 2B, images on left) versus the foci formed 24 hours post-infection (Figure 2B, images on right). Foci were much larger than the input infected cells (Figure 2B bar graph), allowing us to select out by image analysis any input cells which did not lead to infection foci (Materials and Methods).

**Figure 2:**
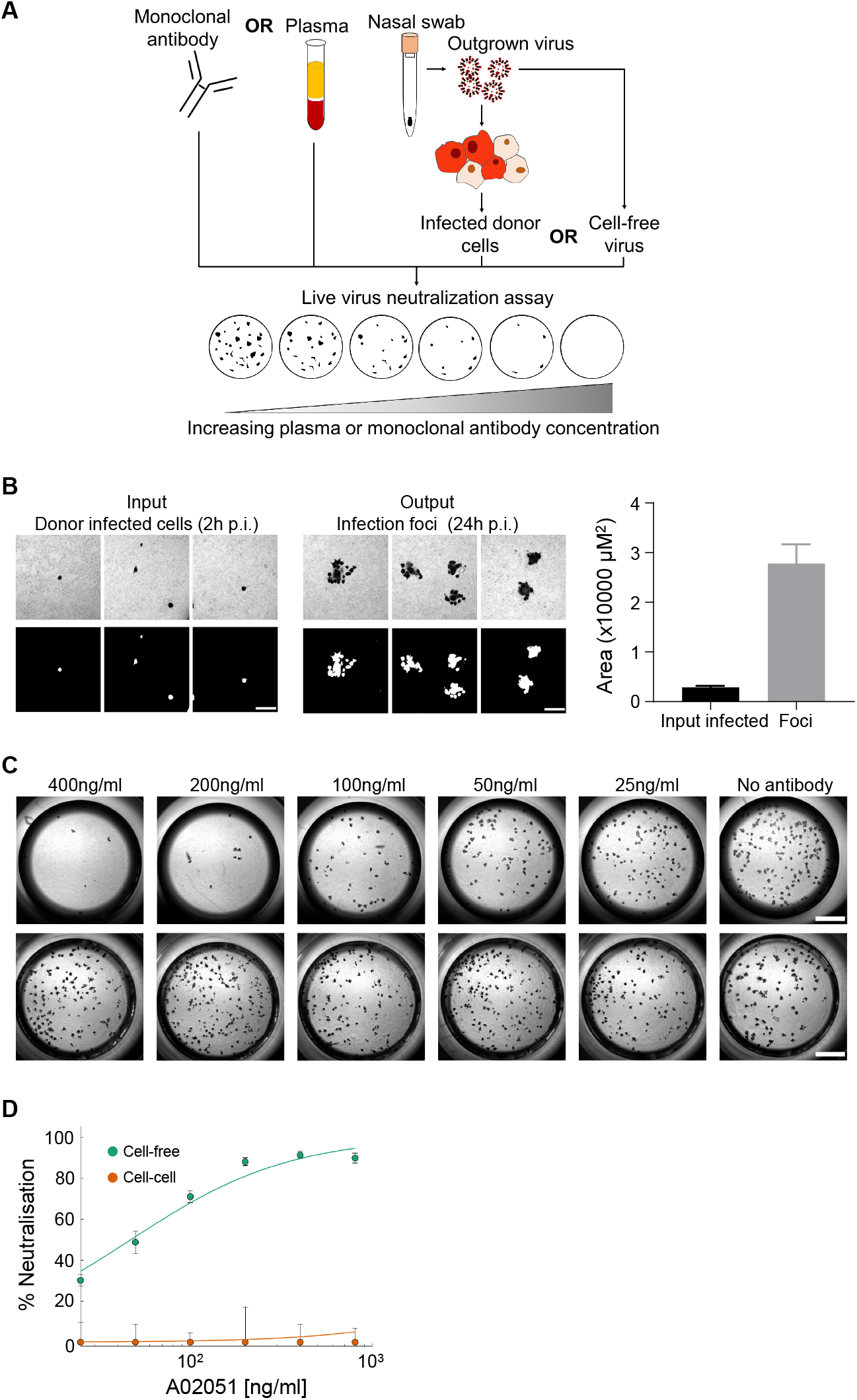
Insensitivity of SARS-CoV-2 cell-to-cell spread to a monoclonal neutralizing antibody. (A) Experimental design. A blood draw and nasopharyngeal/oropharyngeal swab was performed on study participants. Outgrown virus from nasal swabs (D614G or B.1.351) was used to infect H1299-ACE2 cells. Either H1299-ACE2 infected cells or cell-free virus was used as the infection source. Convalescent plasma and a commercial monoclonal antibody A02051 were tested for neutralization capacity using a live virus neutralization assay which detects the number of infection foci. (B) Size differences between input infected cells 2 hours post-addition and foci 24 hours post-infection. Three representative fields of view are shown for the input infected cells (left images) or multi-cell infection foci (middle images) stained with an anti-spike antibody. Bottom image is the masked image showing the labelled infected cell or focus as white. Scale bar is 0.2mm. Bar plot shows automatically detected input infected cell and focus size. Results are from 10 fields of view for input infected cells and 5 fields of view for foci. (C) Focus number in cell-free (top panels) and cell-to-cell infections (bottom panels) in the presence of A02051. Antibody concentration used is above each set of two images. Scale bar is 2mm. (D) Quantified % neutralization. Points are means and SEM from 3 independent experiments.

Visual inspection showed that while cell-free infection was sensitive to neutralization by the mAb, cell-to-cell spread was not (Figure 2C). We normalized the number of foci to the number of foci in the absence of plasma on the same plate to obtain the transmission index (Tx, [4, 20]). This controls for experimental variability in the input virus dose between experiments. We plotted the data for each neutralization experiment as percent neutralization ((1 − *Tx*) × 100%, with values below 0 set to 0. We fitted the data to a sigmoidal function with the antibody concentration required to inhibit 50% of infection (*IC*_50_) as the only free parameter (Materials and Methods). For cell-free infection, *IC*_50_=47.4 ng/ml (95% CI 34.5 to 60.2 ng/ml). In contrast, there was no detectable neutralization by the mAb of cell-to-cell spread (Figure 2D).

### SARS-CoV-2 cell-to-cell spread is not neutralized by convalescent plasma

We next proceeded to test neutralization of virus from plasma of convalescent participants. Five plasma samples were from participants infected during South Africa’s first infection wave, peaking in July 2020 and consisting of viral variants that usually showed the D614G substitution but had none of the defining mutations of B.1.351. Another 6 plasma samples were from participants infected in South Africa’s second wave, peaking in January 2021, when the circulating variant was B.1.351. The infecting virus of most participants was verified by sequencing.

As with the mAb, we mixed the virus or infected cells with serially diluted plasma, then added the mixture to H1299-ACE2 cells and counted infection foci after 24 hours. Plasma samples were tested at serial dilutions ranging from 1:25 to 1:1600. Our previous work showed that neutralization values in different convalescent plasma donors for a given plasma dilution reasonably approximated a normal distribution [20] and we therefore present the mean neutralization across donors (Figure 3A-B). As an alternative, we also present the results from individual donors and their geometric mean (Figure 3C-D). Neutralization for each plasma donor or across donors is represented by the 50% plaque reduction neutralization titer (*PRNT*_50_, [35]), the reciprocal of the plasma dilution required for 50% neutralization. Therefore, a higher number denotes stronger neutralization capacity.

**Figure 3:**
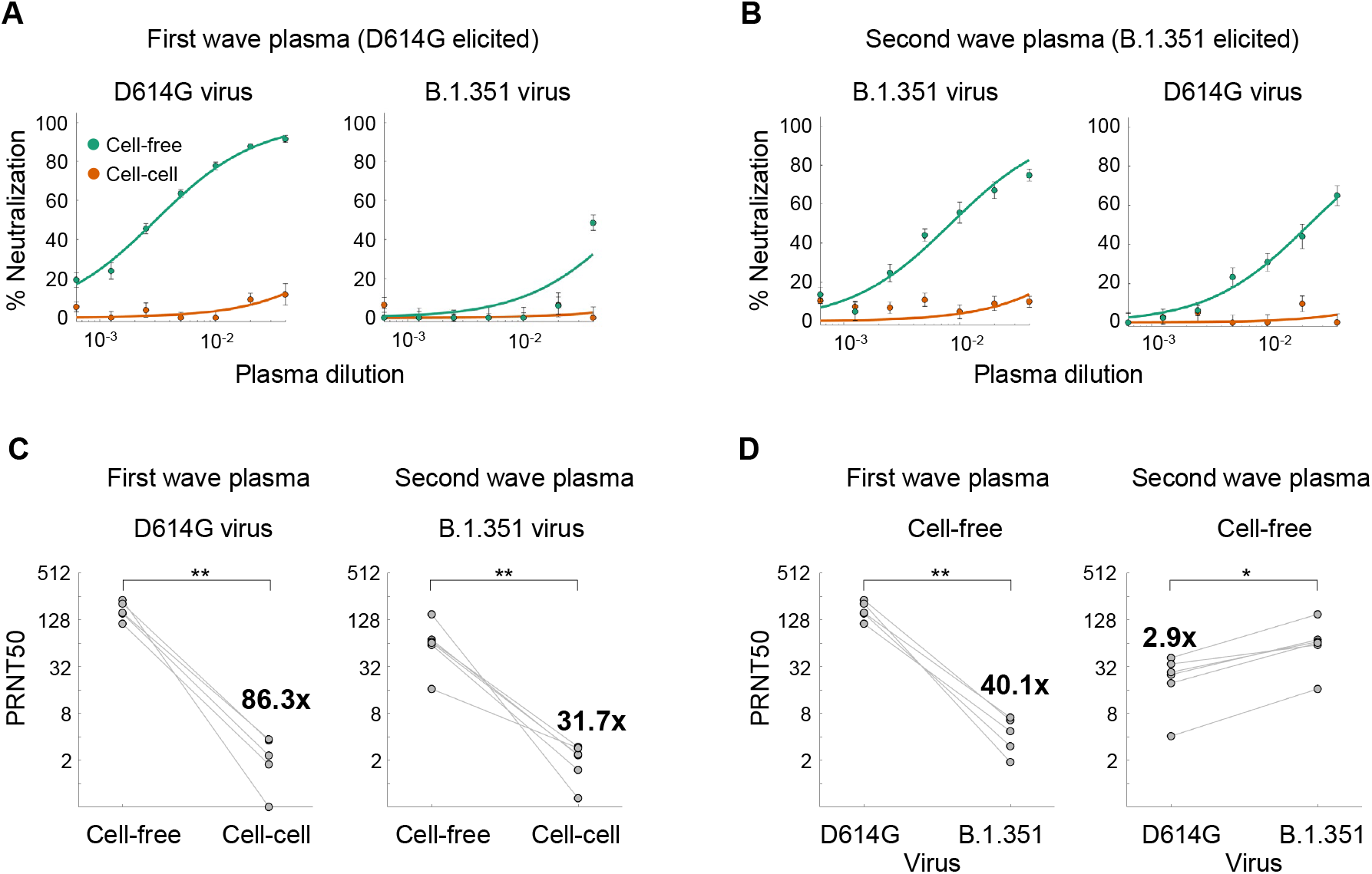
SARS-CoV-2 cell-to-cell spread is insensitive to neutralization by convalescent plasma. (A) Neutralization by first South African infection wave, D614G-elicited plasma of cell-free infection (green) and cell-to-cell spread (orange). Infecting virus was either D614G (left panel) or B.1.351 (right panel). (B) Neutralization by second South African infection wave, B.1.351-elicited plasma of cell-free infection and cell-to-cell spread. Infecting virus was either B.1.351 (left panel) or D614G (right panel). For (A) and (B), points represent mean and SEM of plasma from 5 (A) or 6 (B) study participants, with two independent experiments performed per participant, and curves represent the sigmoidal fits. (C) *PRNT*_50_ of individual participants with D614G elicited plasma when neutralizing D614G cell-free or cell-to-cell infection (left panel) or with B.1.351 elicited plasma when neutralizing B.1.351 cell-free or cell-to-cell infection (right panel). (D) *PRNT*_50_ of individual participants with D614G elicited plasma neutralizing cell-free D614G or B.1.351 infection (left panel) or with B.1.351 elicited plasma neutralizing cell-free D614G or B.1.351 infection (right panel). Fold-change shown in (C) and (D) is the geometric mean of the fold-change per participant between cell-free and cell-to-cell infection (C) or between the homologous virus and the heterologous, cross-neutralized virus (D). p-values are *0.05-0.01 and **0.01-0.001 by the non-parametric Wilcoxon rank sum test.

When cell-free D614G virus was used as the infection source, the *PRNT*_50_ combined across participants was 328.4 (95% CI 290.6-377.4). In contrast, the *PRNT*_50_ for cell-to-cell spread was 3.6 with 95% CI of 2.2-8.7 (Figure 3A, left plot). The value for cell-to-cell spread is an extrapolation, given that our most concentrated plasma dilution was 1:25.

Neutralization capacity of D614G infection elicited convalescent plasma was strongly attenuated with cell-free B.1.351 virus (Figure 3A, right plot), consistent with our previous results in the VeroE6 cell line [20] and the results of others [21, 22, 23, 24]. *PRNT*_50_ across participants was 12.1 (95% CI 6.6-71.3). Cell-to-cell spread reduced *PRNT*_50_ to undetectable (Figure 3B, left plot).

*PRNT*_50_ for neutralization by B.1.351 elicited plasma of B.1.351 cell-free virus was 119.1 (90.4-174.8). Cell-to-cell spread reduced the *PRNT*_50_ to undetectable, given the uncertainty of the fit (Figure 3B, left plot). Cross-neutralization of the cell-free D614G variant with B.1.351 elicited plasma showed a *PRNT*_50_ of 43.7. *PRNT*_50_ for cell-to-cell spread was undetectable.

The analysis of neutralization capacity per participant showed very similar trends to the combined data. The *PRNT*_50_ for neutralization of D614G virus with D614G-elicited plasma ranged between 227.7 to 458.9 (geometric mean 333.9), and dropped 86-fold with cell-to-cell spread (Figure 3C, left plot). The *PRNT*_50_ for neutralization of B.1.351 virus with B.1.351-elicited plasma ranged between 33.0 to 300.2 (geometric mean 119.8), and dropped 32-fold with cell-to-cell spread (Figure 3C, right plot).

Comparison of cell-free infection in H1299-ACE2 cells showed that B.1.351 was poorly cross-neutralized with D614G-elicited plasma. *PRNT*_50_ values for cross-neutralization ranged from 3.8 to 14.2, with a geometric mean of 8.3, a 40-fold drop. In contrast, B.1.351-elicited plasma did not show as large of a decrease in neutralization capacity when neutralizing D614G virus (range: 8.2-82.8, geometric mean 41.3) with a fold-change of about 3.

## Discussion

We have shown that SARS-CoV-2 infection originating in infected cells is insensitive to neutralizing antibodies. The importance of these results lies in the fact that in natural immunity, the antibody response requires several weeks to reach its peak [36]. Given the rapid infection cycle of SARS-CoV-2 [34], neutralization in natural infection very likely occurs after cells have been infected. If active infection is still present, neutralization may not be effective to the same degree in all infected cell types and anatomical compartments because of cell-to-cell spread of SARS-CoV-2. T cell immunity [37] to SARS-CoV-2 may be required clear compartments refractory to neutralization.

Unlike natural infection, the antibodies elicited by a vaccine should be present before the exposure to the virus. A vaccine eliciting a strong neutralizing antibody response which lasts for months could potentially clear the cell-free infection before it infects cells or cells capable of cell-to-cell spread. If lower neutralization capacity is elicited by the vaccine, some cells could be infected, since any vaccine mediated T cell immunity would target infected cells, not the transmitted virus, and therefore some cellular infection would need to occur. Therefore, syncytia formation could potentially happen.

A caveat to our results is that we used a human cancer cell line, and modified it to express ACE-2. However, syncytia are a common feature of SARS-CoV-2 lung pathology [11, 12, 13], showing that the cell fusion mechanism of cell-to-cell spread operates in this environment. We used infected cells after they already expressed the SARS-CoV-2 spike protein on the cell surface, providing a target for neutralization. Despite this, there was no appreciable neutralization.

Both the earlier D614G variant and the B.1.351 variant showed similar insensitivity of cell-to-cell spread in D614G and B.1.351 elicited plasma, indicating that the insensitivity of cell-to-cell spread to neutralization is not specific to the infecting variant or the elicited neutralizing antibodies. We have also verified previous observations about the drop in B.1.351 neutralization by non-B.1.351 elicited plasma in the H1299-ACE2 human lung cell system, as well as the cross-neutralization of D614G by B.1.351-elicited plasma. Interestingly, in these experiments, neutralization of B.1.351 by its matched plasma was weaker than neutralization of D614G by its matched plasma. Despite the overall weaker neutralization capacity, the B.1.351-elicited plasma could still effectively cross-neutralize, showing that cross-neutralization is not necessarily linked to absolute neutralization capacity.

Like with other viruses, cell-to-cell spread of SARS-CoV-2 may prove to play a role in pathology and possibly persistence. Future vaccine and therapeutic strategies should consider approaches to minimize this mode of transmission.

## Material and Methods

### Ethical statement

Nasopharyngeal/oropharyngeal swab samples and plasma samples were obtained from 11 hospitalized adults with PCR-confirmed SARS-CoV-2 infection enrolled in a prospective cohort study approved by the Biomedical Research Ethics Committee (BREC) at the University of KwaZulu-Natal (reference BREC/00001275/2020). The D614G and B.1.351 isolates were outgrown from a swab taken as part of the cohort study (D614G) or from a residual nasopharyngeal/oropharyngeal sample used for routine SARS-CoV-2 diagnostic testing through our SARS-CoV-2 genomic surveillance program (BREC approval reference BREC/00001510/2020).

### Data and code availability statement

Sequence data that support the findings of this study have been deposited in GISAID.

### Cells

Vero E6 cells (ATCC CRL-1586, obtained from Cellonex in South Africa) were propagated in complete DMEM with 10% fetal bovine serum (Hylone) containing 1% each of HEPES, sodium pyruvate, L-glutamine, and non-essential amino acids (Sigma-Aldrich). Cells were passaged every 3-4 days. H1299 and H1299 subclones were propagated in complete RPMI with 10% fetal bovine serum containing 1% each of HEPES, sodium pyruvate, L-glutamine, and non-essential amino acids and and passaged every second day. HEK-293 (ATCC CRL-1573) cells were propagated in complete DMEM with 10% fetal bovine serum containing 1% each of HEPES, sodium pyruvate, L-glutamine, and non-essential amino acids and and passaged every second day.

### Single cell cloning to create H1299-ACE2 clones

The H1299-H2AZ clone with nuclear labelled YFP was constructed to overexpress ACE2 as follows: VSVG-pseudotyped lentivirus containing the human ACE2 was generated by co-transfecting 293T cells with the pHAGE2-EF1alnt-ACE2-WT plasmid along with the lentiviral helper plasmids HDM-VSVG, HDM-Hgpm2, HDM-tat1b and pRC-CMV-Rev1b using TransIT-LT1 (Mirus) transfection reagent. Supernatant containing the lentivirus was harvested two days after infection, filtered through a 0.45*μ*m filter (Corning) and used to spinfect H1299-H2AZ at 1000 rcf for 2 hours at room temperature in the presence of 5 *μ*g/mL polybrene (Sigma-Aldrich). ACE-2 transduced H1299-H2AZ cells were then sub-cloned at the single cell density in 384-well plates (Greiner Bio-One) in conditioned media derived from H1299 confluent cells. After 3 weeks, wells that had good growth of cell were trypsinized (Sigma-Aldrich) and plated in two replicate 96-well plates (Corning), where the first plate was used to determine infectivity and the second was stock. The first plate was screened for the fraction of mCherry positive cells per cell clone upon infection with SARS-CoV-2 mCherry expressing spike pseudotyped lentiviral vector 1610-pHAGE2/EF1a Int-mCherry3-W produced by transfecting as above. Screening was performed using a Metamorph-controlled (Molecular Devices, Sunnyvale, CA) Nikon TiE motorized microscope (Nikon Corporation, Tokyo, Japan) with a 20x, 0.75 NA phase objective, 561 laser line, and 607 nm emission filter (Semrock, Rochester, NY). Images were captured using an 888 EMCCD camera (Andor). Temperature (37°C), humidity and CO2 (5%) were controlled using an environmental chamber (OKO Labs, Naples, Italy). The three clones with the highest fraction of mCherry expression was expanded from the stock plate and denoted H1299-C1, H1299-C7 and H1299-E3.

### Time-lapse microscopy

To produce infected donor cells, the three subclones of H1299 (C1, C7 and E3) were seeded at 4 × 10^5^ cells/ml 1 day pre-infection. D614G virus was then added to the cells and cells were incubated for 16 hours. On the same day as the donor cell infection, target cells (H1299-C1, -C7 and -E3) were seeded at 20 × 10^4^ cells/ml, in a total of 3ml, into duplicate wells of a fibronectin (Sigma-Aldrich) coated 6-well optical plate (MaTek). 18 hours post target cell seeding and donor cell infection, infected donor cells were trypsinized, resuspended at 20 × 10^4^ cells/ml and added to the matched target cells at a 1:20 dilution. For time-lapse microscopy, donor-target infections were imaged in tandem using a Metamorph-controlled Nikon TiE motorized microscope with environmental control as above in a biosafety level 3 facility. Excitation source was 488 laser line and emission was detected through a Semrock Brightline quad band 440–40 /521-21/607-34/700-45 nm filter. For each well, four fields of view were captured every 10 minutes for a total of 46 hours.

### Image analysis of multinucleated H1299-ACE2 cells in time lapse microscopy images

Timelapse microscopy images were analysed using custom Matlab script. Fluorescently labelled nuclei in the YFP channel were used to quantify the nucleation state of cell lines (Figure S2). Background illumination signal in the YFP fluorescent channel was removed by subtracting 10% of maximum fluorescence from the image. Fluorescent signal from YFP labelled nuclei was then used to generate a binary mask and select individual nucleus objects in each image. Each nucleus object was categorized as multi-nucleate or uni-nucleate based on the maximum area of a single nucleus. The maximum area, of a single nucleus, for each cell line was determined by identifying the largest single nucleus (by pixel area) in the uninfected controls. Each nucleus object was also assigned a size value relative to the average size of a single nucleus for a cell line. The average nucleus size (by pixel area), per cell line, was calculated from single nuclei in uninfected controls. Data generated in this way was graphed in R using the ggplot2 package. Data from all three cell lines was used to calculate mean and standard deviation of the percentage of nuclei that were multi-nucleate at each timepoint, as well as the number of nuclei at each timepoint for infected and uninfected cells.

### Isolation of plasma from blood

Plasma was separated from EDTA-anticoagulated blood by centrifugation at 500 rcf for 10 minutes and stored at −80°C. Aliquots of plasma samples were heat-inactivated at 56°C for 30 minutes, and clarified by centrifugation at 10,000 rcf for 5 minutes, where the clear middle layer was used for experiments. Inactivated plasma was stored in single use aliquots to prevent freeze-thaw cycles.

### LVNA using focus forming assay

To quantify neutralization of virus either in cell-free or cell-to-cell infections, H1299-C7 cells were plated in an 96-well plate (Corning) at 30,000 target cells per well 1 day pre-infection. Importantly, before infection approximately 5ml of sterile water was added between wells to prevent more drying of wells at the edge of the plate which we have observed to cause edge effects. For the neutralization, plasma was serially diluted two-fold from 1:25 to 1:1600, where this is the concentration during the virus-plasma/ infected donor cell-plasma incubation step before addition to target cells. Donor cells were prepared by infecting cells 1 day pre-infection with either D614G or B.1.351 virus. Briefly, virus was added to donor cells seeded at 4 × 10^5^ cells/ml and incubated for 16 hours. These infected donor cells were trypsinized, spun down at 300 rcf for 3 minutes and resuspended at 20 × 10^4^ cells/ml in fresh growth medium – calibrated to result in approximately 100 focus-forming units (FFU) per microwell in the target cell plate. Before addition to target cells, the cell suspension was allowed to stand for 5 minutes to separate cell syncytia from the single infected cells to be used as the infection source. The cell suspension excluding precipitated cell syncytia and debris was validated using a microscope to contain only single cells and used for the cell-to-cell infections. Virus stocks (for cell-free infections) were used at approximately 100 focus-forming units (FFU) per microwell. Both donor cells and virus were added to diluted plasma. The antibody-virus or antibody-donor cell mixtures were incubated for 1 hour at 37°C, 5% CO2. Target cells were infected with 100*μ*L (1:2 of final volume) of the antibody-virus or antibody-donor cell mixtures for one hour, to allow adsorption of virus. Subsequently, 100*μ*L of a 1x RPMI 1640 (Sigma-Aldrich R6504), 1.5% carboxymethylcellulose (Sigma-Aldrich C4888) overlay was added to the wells without removing the inoculum. Cells were fixed at 24 hours post-infection using 4% paraformaldehyde (Sigma-Aldrich) for 20 minutes. For staining of foci, a rabbit anti-spike monoclonal antibody (mAb BS-R2B12, GenScript A02058) was used at 0.5*μ*g/mL as the primary detection antibody. Antibody was resuspended in a permiabilization buffer containing 0.1% saponin (Sigma-Aldrich), 0.1% BSA (Sigma-Aldrich), and 0.05% tween (Sigma-Aldrich) in PBS. Plates were incubated with primary antibody overnight at 4°C, then washed with wash buffer containing 0.05% tween in PBS. Secondary goat anti-rabbit horseradish peroxidase (Abcam ab205718) was added at 1 *μ*g/mL and incubated for 2 hours at room temperature with shaking. The TrueBlue peroxidase substrate (SeraCare 5510-0030) was then added at 50*μ*L per well and incubated for 20 minutes at room temperature. Plates were then dried for 2 hours and imaged using a Metamorph-controlled Nikon TiE motorized microscope with a 2x objective (Nikon). Automated image analysis was performed using a Matlab2019b (Mathworks) custom script, where focus detection was automated and did not involve user curation. Image segmentation steps were stretching the image from minimum to maximum intensity, local Laplacian filtering, image complementation, thresholding and binarization.

### Statistics and fitting

All statistics and fitting were performed using Matlab2019b. Neutralization data was fit to

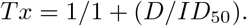

Here Tx is the number of foci normalized to the number of foci in the absence of plasma on the same plate at dilution D. To visualize the data, we used percent neutralization, calculated as (1-Tx) × 100%. Negative values (Tx>1, enhancement) were represented as 0% neutralization. Fit to a normal distribution used Matlab2019b function normplot, which compared the distribution of the Tx data to the normal distribution (see https://www.mathworks.com/help/stats/normplot.html).

### ACE2 antibody staining

H1299-C7 cells were seeded at 4 × 10^5^ cells/ml, in a total of 3 ml, into wells of a 6-well optical plate (Matek), 1 day prior to staining. For staining with ACE2, cells were washed twice with DPBS and then fixed with 2 mL/well 4% paraformaldehyde (Sigma-Aldrich) for 30 minutes. The paraformaldehyde was removed and cells were washed three times with DPBS. Primary ACE-2 antibody (Abcam ab272690) was added to cells at a concentration of 2.5 *μ*g/mL in an antibody staining solution composed of 1% Bovine Serum Albumin (Sigman-Aldrich) in DPBST. Cells were incubated with ACE2 antibody overnight at 4°C with rocking. Cells were then washed three times with DPBS and incubated with secondary antibody, goat anti-rabbit IgG Alexa Fluor 555 (Abcam ab150078) at a concentration of 4 *μ*g/mL in an antibody staining solution made up of 1% Bovine Serum Albumin (Sigman-Aldrich) in DPBST. The nuclear DAPI stain (Sigma-Aldrich) was used to counter-stain nuclei and added to the cells at a 0.5 *μ*g/mL. For imaging, a Metamorph-controlled Nikon TiE motorized microscope with a 60x oil immersion objective, 0.75 NA phase objective was used. Excitation source was 515 (Alexa Fluor 555) and 405 (DAPI) laser lines and emission was detected through a Semrock Brightline quad band 440–40 /521-21/607-34/700-45 nm filter. Images were captured using an 888 EMCCD camera (Andor).

## Acknowledgements

This work was supported by the Bill and Melinda Gates Investment INV-018944 (AS) and The Africa Health Research Institute (AHRI) internal seed funding for COVID-19 research (LJ).

## Author Contributions

Conceived study and designed experiments: AS and LJ. Performed experiments: LJ, HR, SHH, SC, YG. Analyzed and Interpreted data: AS, HR, and LJ. Cohort set-up and management: FK, KK, MB, YG, BIG, MYM, SC, AS, WH, AS. Sequencing and analysis: TdO, JG, HT. AS, HR, and LJ prepared the manuscript.

## COMMIT-KZN Team

Moherndran Archary^36^, Kaylesh J. Dullabh^37^, Philip Goulder^1,38^, Sashin Harilall^1^, Guy Harling^1,39^, Rohen Harrichandparsad^40^, Kobus Herbst^1,41^, Prakash Jeena^36^, Thandeka Khoza^1^, Nigel Klein^1, 42^, Henrik Kløverpris^1, 5, 43^, Alasdair Leslie^1, 5^, Rajhmun Madansein^37^, Mohlopheni Marakalala^1, 5^, Matilda Mazibuko^1^, Mosa Moshabela^44^, Ntombifuthi Mthabela^1^, Kogie Naidoo^6^, Zaza Ndhlovu^1, 7^, Thumbi Ndung’u^1, 5, 10, 45^, Kennedy Nyamande^46^, Nesri Padayatchi^6^, Vinod Patel^47^, Dirhona Ramjit^1^, Hylton Rodel^1,5^, Theresa Smit^1^, Adrie Steyn^1,48^, Emily Wong^1,48^.

^36^Department of Paediatrics and Child Health, University of KwaZulu-Natal, Durban, South Africa. ^37^Department of Cardiothoracic Surgery, University of KwaZulu-Natal, Durban, South Africa. ^1^Africa Health Research Institute, Durban, South Africa. 3^8^Department of Paediatrics, Oxford, UK. ^39^Institute for Global Health, University College London, UK. ^40^Department of Neurosurgery, University of KwaZulu-Natal, Durban, South Africa. ^41^South African Population Research Infrastructure Network, Durban South Africa. ^42^Institute of Child Health, University College London, UK. 5Division of Infection and Immunity, University College London, London, UK. ^43^Department of Immunology and Microbiology, University of Copenhagen, Copenhagen, Denmark. ^44^College of Health Sciences, University of KwaZulu-Natal, Durban, South Africa. ^6^Centre for the AIDS Programme of Research in South Africa, Durban,South Africa. ^7^Ragon Institute of MGH, MIT and Harvard, Boston, USA. ^45^HIV Pathogenesis Programme, The Doris Duke Medical Research Institute, University of KwaZulu-Natal, Durban, South Africa. ^10^Max Planck Institute for Infection Biology, Berlin, Germany. ^46^Department of Pulmonology and Critical Care, University of KwaZulu-Natal, Durban, South Africa. ^47^Department of Neurology, University of KwaZulu-Natal, Durban, South Africa. ^48^Division of Infectious Diseases, University of Alabama at Birmingham.

---

**Figure S1:**
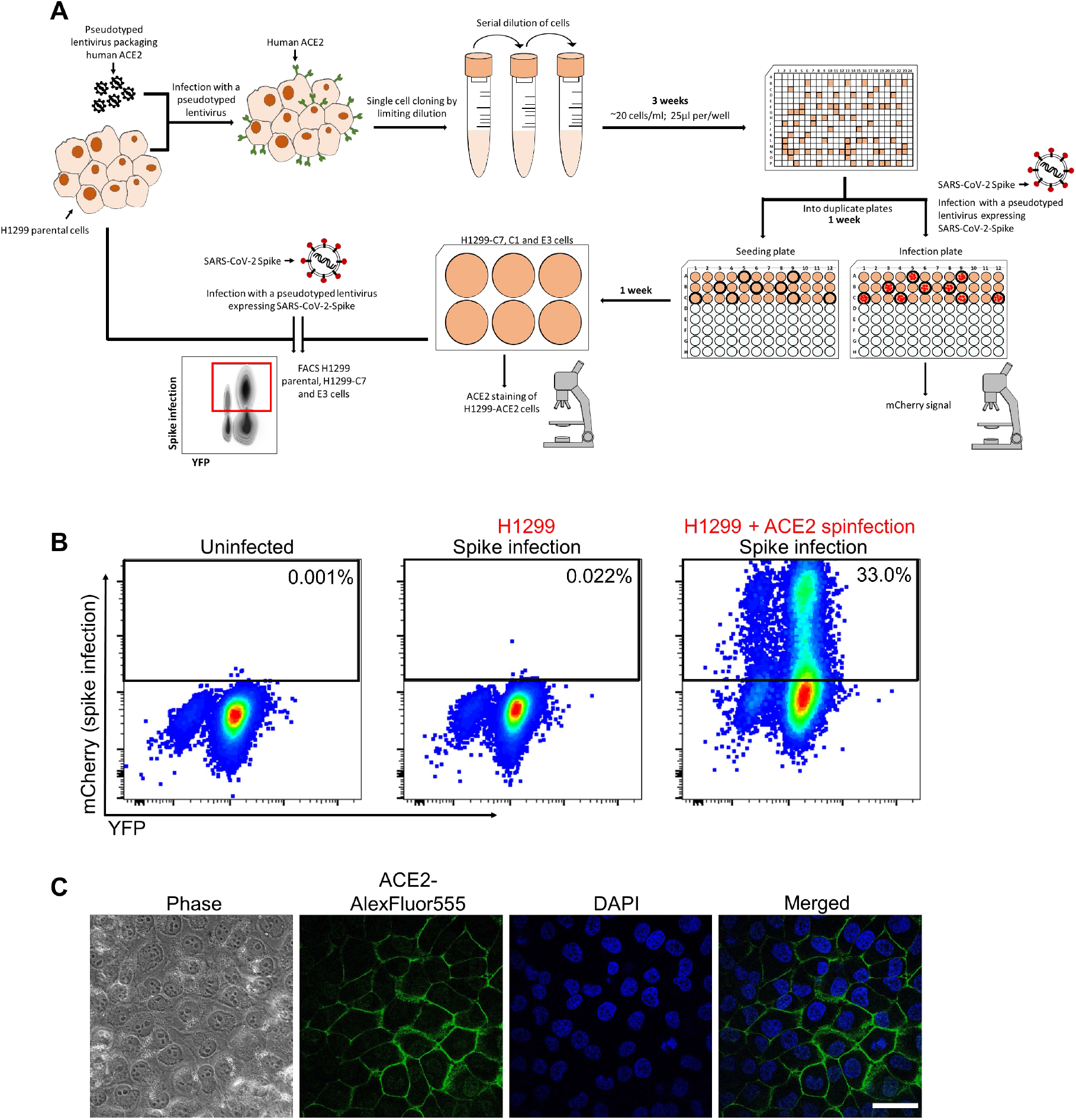
Generation of the H1299-C1, -C7 and -E3 clonal cell lines. (A) The H1299 epithelial cell line with YFP labelled H2AZ was spinfected with the pHAGE2-EF1a-Int-ACE2 lentivector. Cells were single cell cloned by limiting dilution in a 384 well plate. Clones were expanded into duplicate 96-well plates, where one plate was used for seeding and the other plate was used to select infectable clones based on mCherry signal from infection with SARS-CoV-2 mCherry expressing spike pseudotyped lentivirus. (B) Flow cytometry plots of SARS-CoV-2 mCherry expressing spike pseudotyped lentivirus infection in H1299-C7 cells. (C) Images of H1299-C7 cells stained with ACE2 and DAPI. Scale bar is 50*μ*m.

**Figure S2:**
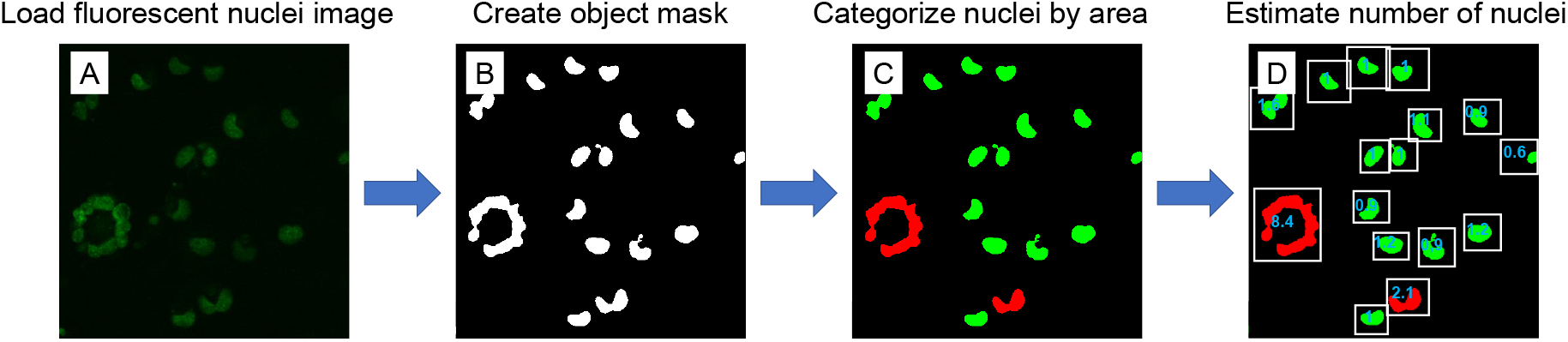
Schematic overview of multinucleate image analysis. (A) Images of fluorescently labelled nuclei were loaded into Matlab and a (B) binary mask was generated to separate individual nuclei objects. (C) An object was classified as multinucleate (red) if the object’s area was larger than the maximum determined size for a single nucleus, and uni-nucleate (green) otherwise. (D) The number of nuclei in each object was then determined by comparing it to the average area of a single nucleus.

**Figure S3:**
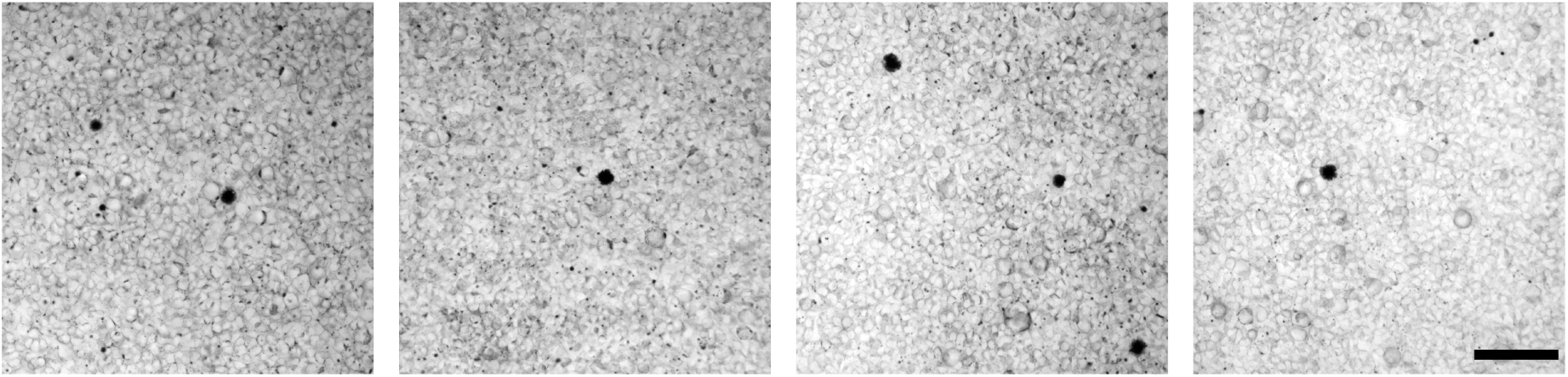
Surface expression of spike on input infected donor cells. Cells were infected by cell-free virus for 16 hours before they were used as the infecting donor cells. Infected donor cells were trypsinized and added to target cells. Donor-target cell mixture was fixed and stained with rabbit anti-spike monoclonal antibody (A02058) in staining solution without saponin, therefore staining any spike on the cell surface only. Each image represents a different field of view of the donor-target cell mixture. Scale bar is 200*μ*m.

**Figure S4:**
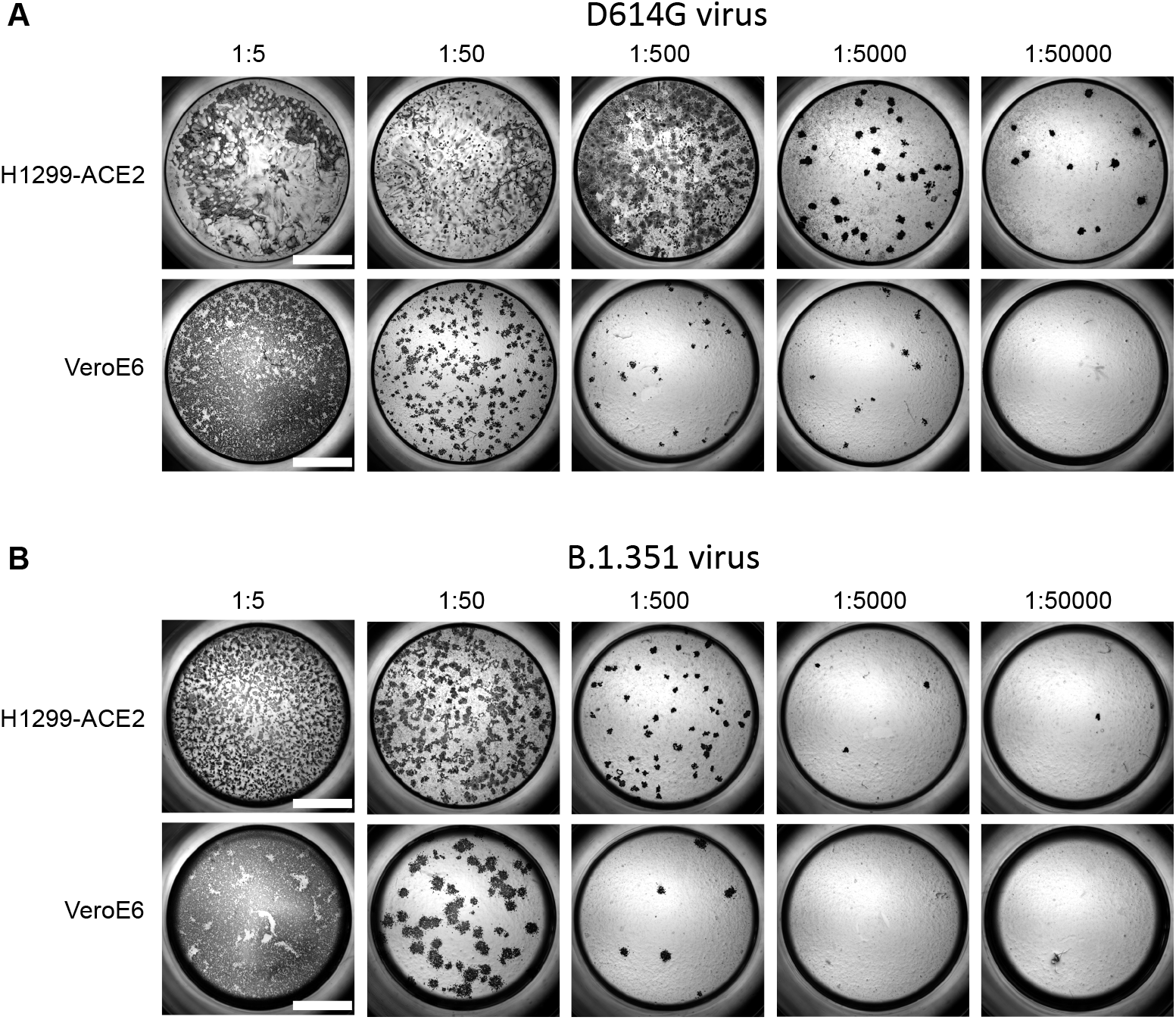
Comparision of SARS-CoV-2 infection in H1299-ACE2 and VeroE6 cells. Both H1299-ACE2 and VeroE6 cells were infected with the same viral stock in the same experiment with D614G virus (A) or B.1.351 virus (B) at different dilutions and focus forming assay was performed. Scale bar is 2mm.

## References

[1] Q. Sattentau. Avoiding the void: cell-to-cell spread of human viruses. Nat Rev Microbiol, 6(11): 815–26 2008. ISSN 1740-1534 (Electronic) 1740-1526 (Linking). doi: 10.1038/nrmicro1972. URL https://www.ncbi.nlm.nih.gov/pubmed/18923409.

[2] Clare Jolly, Kirk Kashefi, Michael Hollinshead, and Quentin J Sattentau. Hiv-1 cell to cell transfer across an env-induced, actin-dependent synapse. The Journal of experimental medicine, 199(2): 283–293 2004.

[3] Stefanie Sowinski, Clare Jolly, Otto Berninghausen, Marco A Purbhoo, Anne Chauveau, Karsten Köhler, Stephane Oddos, Philipp Eissmann, Frances M Brodsky, Colin Hopkins, et al. Membrane nanotubes physically connect t cells over long distances presenting a novel route for hiv-1 transmission. Nature cell biology, 10(2):211–219, 2008.

[4] A. Sigal, J. T. Kim, A. B. Balazs, E. Dekel, A. Mayo, R. Milo, and D. Baltimore. Cell-to-cell spread of HIV permits ongoing replication despite antiretroviral therapy. Nature, 477(7362):95–8, 2011. ISSN 1476-4687 (Electronic) 0028-0836 (Linking). doi: 10.1038/nature10347. URL https://www.ncbi.nlm.nih.gov/pubmed/21849975.

[5] Irene A Abela, Livia Berlinger, Merle Schanz, Lucy Reynell, Huldrych F Günthard, Peter Rusert, and Alexandra Trkola. Cell-cell transmission enables hiv-1 to evade inhibition by potent cd4bs directed antibodies. PLoS Pathog, 8(4):e1002634, 2012.

[6] Torben Schiffner, Quentin J Sattentau, and Christopher JA Duncan. Cell-to-cell spread of hiv-1 and evasion of neutralizing antibodies. Vaccine, 31(49):5789–5797, 2013.

[7] A. Moyano, G. Lustig, H. E. Rodel, T. Antal, and A. Sigal. Interference with HIV infection of the first cell is essential for viral clearance at sub-optimal levels of drug inhibition. PLoS Comput Biol, 16(2):e1007482, 2020. ISSN 1553-7358 (Electronic) 1553-734X (Linking). doi: 10.1371/journal.pcbi.1007482. URL https://www.ncbi.nlm.nih.gov/pubmed/32017770.

[8] Mehdi Bouhaddou, Danish Memon, Bjoern Meyer, Kris M White, Veronica V Rezelj, Miguel Correa Marrero, Benjamin J Polacco, James E Melnyk, Svenja Ulferts, Robyn M Kaake, et al. The global phosphorylation landscape of sars-cov-2 infection. Cell, 182(3):685–712, 2020.

[9] J. Buchrieser, J. Dufloo, M. Hubert, B. Monel, D. Planas, M. M. Rajah, C. Planchais, F. Porrot, F. Guivel-Benhassine, S. Van der Werf, N. Casartelli, H. Mouquet, T. Bruel, and O. Schwartz. Syncytia formation by sars-cov-2-infected cells. EMBO J, 39(23):e106267, 2020. ISSN 1460-2075 (Electronic) 0261-4189 (Linking). doi: 10.15252/embj.2020106267. URL https://www.ncbi.nlm.nih.gov/pubmed/33051876.

[10] M. Hoffmann, H. Kleine-Weber, and S. Pohlmann. A multibasic cleavage site in the spike protein of sars-cov-2 is essential for infection of human lung cells. Mol Cell, 78(4):779–784 e5, 2020. ISSN 1097-4164 (Electronic) 1097-2765 (Linking). doi: 10.1016/j.molcel.2020.04.022. URL https://www.ncbi.nlm.nih.gov/pubmed/32362314.

[11] Luca Braga, Hashim Ali, Ilaria Secco, Elena Chiavacci, Guilherme Neves, Daniel Goldhill, Rebecca Penn, Jose M Jimenez-Guardeño, Ana M Ortega-Prieto, Rossana Bussani, et al. Drugs that inhibit tmem16 proteins block sars-cov-2 spike-induced syncytia. Nature, pages 1–6, 2021.

[12] R. Bussani, E. Schneider, L. Zentilin, C. Collesi, H. Ali, L. Braga, M. C. Volpe, A. Colliva, F. Zan-conati, G. Berlot, F. Silvestri, S. Zacchigna, and M. Giacca. Persistence of viral rna, pneumocyte syncytia and thrombosis are hallmarks of advanced covid-19 pathology. EBioMedicine, 61:103104, 2020. ISSN 2352-3964 (Electronic) 2352-3964 (Linking). doi: 10.1016/j.ebiom.2020.103104. URL https://www.ncbi.nlm.nih.gov/pubmed/33158808.

[13] Liangyu Lin, Qing Li, Ying Wang, and Yufang Shi. Syncytia formation during sars-cov-2 lung infection: a disastrous unity to eliminate lymphocytes. Cell Death & Differentiation, pages 1–3, 2021.

[14] D. S. Khoury, D. Cromer, A. Reynaldi, T. E. Schlub, A. K. Wheatley, J. A. Juno, K. Subbarao, S. J. Kent, J. A. Triccas, and M. P. Davenport. Neutralizing antibody levels are highly predictive of immune protection from symptomatic sars-cov-2 infection. Nat Med, 2021. ISSN 1546-170X (Electronic) 1078-8956 (Linking). doi: 10.1038/s41591-021-01377-8. URL https://www.ncbi.nlm.nih.gov/pubmed/34002089.

[15] Kristen A. Earle, Donna M. Ambrosino, Andrew Fiore-Gartland, David Goldblatt, Peter B. Gilbert, George R. Siber, Peter Dull, and Stanley A. Plotkin. Evidence for antibody as a protective correlate for covid-19 vaccines. medRxiv, 2021. doi: 10.1101/2021.03.17.20200246. URL https://www.medrxiv.org/content/early/2021/03/20/2021.03.17.20200246.

[16] C. O. Barnes, C. A. Jette, M. E. Abernathy, K. A. Dam, S. R. Esswein, H. B. Gristick, A. G. Malyutin, N. G. Sharaf, K. E. Huey-Tubman, Y. E. Lee, D. F. Robbiani, M. C. Nussenzweig, Jr. West, A. P., and P. J. Bjorkman. SARS-CoV-2 neutralizing antibody structures inform therapeutic strategies. Nature, 588(7839):682–687, 2020. ISSN 1476-4687 (Electronic) 0028-0836 (Linking). doi: 10.1038/s41586-020-2852-1. URL https://www.ncbi.nlm.nih.gov/pubmed/33045718.

[17] L. Piccoli, Y. J. Park, M. A. Tortorici, N. Czudnochowski, A. C. Walls, M. Beltramello, C. Silacci-Fregni, D. Pinto, L. E. Rosen, J. E. Bowen, O. J. Acton, S. Jaconi, B. Guarino, A. Minola, F. Zatta, N. Sprugasci, J. Bassi, A. Peter, A. De Marco, J. C. Nix, F. Mele, S. Jovic, B. F. Rodriguez, S. V. Gupta, F. Jin, G. Piumatti, G. Lo Presti, A. F. Pellanda, M. Biggiogero, M. Tarkowski, M. S. Pizzuto, E. Cameroni, C. Havenar-Daughton, M. Smithey, D. Hong, V. Lepori, E. Albanese, A. Ceschi, E. Bernasconi, L. Elzi, P. Ferrari, C. Garzoni, A. Riva, G. Snell, F. Sallusto, K. Fink, H. W. Virgin, A. Lanzavecchia, D. Corti, and D. Veesler. Mapping neutralizing and immunodominant sites on the SARS-CoV-2 spike receptor-binding domain by structure-guided high-resolution serology. Cell, 183(4):1024–1042 e21, 2020. ISSN 1097-4172 (Electronic) 0092-8674 (Linking). doi: 10.1016/j.cell.2020.09.037. URL https://www.ncbi.nlm.nih.gov/pubmed/32991844.

[18] M. McCallum, A. De Marco, F. A. Lempp, M. A. Tortorici, D. Pinto, A. C. Walls, M. Beltramello, A. Chen, Z. Liu, F. Zatta, S. Zepeda, J. di Iulio, J. E. Bowen, M. Montiel-Ruiz, J. Zhou, L. E. Rosen, S. Bianchi, B. Guarino, C. S. Fregni, R. Abdelnabi, S. C. Foo, P. W. Rothlauf, L. M. Bloyet, F. Benigni, E. Cameroni, J. Neyts, A. Riva, G. Snell, A. Telenti, S. P. J. Whelan, H. W. Virgin, D. Corti, M. S. Pizzuto, and D. Veesler. N-terminal domain antigenic mapping reveals a site of vulnerability for sars-cov-2. Cell, 184(9):2332–2347 e16, 2021. ISSN 1097-4172 (Electronic) 0092-8674 (Linking). doi: 10.1016/j.cell.2021.03.028. URL https://www.ncbi.nlm.nih.gov/pubmed/33761326.

[19] H. Tegally, E. Wilkinson, M. Giovanetti, A. Iranzadeh, V. Fonseca, J. Giandhari, D. Doolabh, S. Pillay, E. J. San, N. Msomi, K. Mlisana, A. von Gottberg, S. Walaza, M. Allam, A. Ismail, T. Mohale, A. J. Glass, S. Engelbrecht, G. Van Zyl, W. Preiser, F. Petruccione, A. Sigal, D. Hardie, G. Marais, M. Hsiao, S. Korsman, M. A. Davies, L. Tyers, I. Mudau, D. York, C. Maslo, D. Goedhals, S. Abrahams, O. Laguda-Akingba, A. Alisoltani-Dehkordi, A. Godzik, C. K. Wibmer, B. T. Sewell, J. Lourenco, L. C. J. Alcantara, S. L. Kosakovsky Pond, S. Weaver, D. Martin, R. J. Lessells, J. N. Bhiman, C. Williamson, and T. de Oliveira. Emergence of a SARS-CoV-2 variant of concern with mutations in spike glycoprotein. Nature, 2021. ISSN 1476-4687 (Electronic) 0028-0836 (Linking). doi: 10.1038/s41586-021-03402-9. URL https://www.ncbi.nlm.nih.gov/pubmed/33690265.

[20] S. Cele, I. Gazy, L. Jackson, S. H. Hwa, H. Tegally, G. Lustig, J. Giandhari, S. Pillay, E. Wilkinson, Y. Naidoo, F. Karim, Y. Ganga, K. Khan, M. Bernstein, A. B. Balazs, B. I. Gosnell, W. Hanekom, M. S. Moosa, Africa Network for Genomic Surveillance in South, Commit-Kzn Team, R. J. Lessells, T. de Oliveira, and A. Sigal. Escape of sars-cov-2 501y.v2 from neutralization by convalescent plasma. Nature, 593(7857):142–146, 2021. ISSN 1476-4687 (Electronic) 0028-0836 (Linking). doi: 10.1038/s41586-021-03471-w. URL https://www.ncbi.nlm.nih.gov/pubmed/33780970.

[21] C. K. Wibmer, F. Ayres, T. Hermanus, M. Madzivhandila, P. Kgagudi, B. Oosthuysen, B. E. Lambson, T. de Oliveira, M. Vermeulen, K. van der Berg, T. Rossouw, M. Boswell, V. Ueckermann, S. Meiring, A. von Gottberg, C. Cohen, L. Morris, J. N. Bhiman, and P. L. Moore. Sars-cov-2 501y.v2 escapes neutralization by south african covid-19 donor plasma. Nat Med, 27(4):622–625, 2021. ISSN 1546-170X (Electronic) 1078-8956 (Linking). doi: 10.1038/s41591-021-01285-x. URL https://www.ncbi.nlm.nih.gov/pubmed/33654292.

[22] D. Planas, T. Bruel, L. Grzelak, F. Guivel-Benhassine, I. Staropoli, F. Porrot, C. Planchais, J. Buchrieser, M. M. Rajah, E. Bishop, M. Albert, F. Donati, M. Prot, S. Behillil, V. Enouf, M. Maquart, M. Smati-Lafarge, E. Varon, F. Schortgen, L. Yahyaoui, M. Gonzalez, J. De Seze, H. Pere, D. Veyer, A. Seve, E. Simon-Loriere, S. Fafi-Kremer, K. Stefic, H. Mouquet, L. Hocqueloux, S. van der Werf, T. Prazuck, and O. Schwartz. Sensitivity of infectious sars-cov-2 b.1.1.7 and b.1.351 variants to neutralizing antibodies. Nat Med, 27(5):917–924, 2021. ISSN 1546-170X (Electronic) 1078-8956 (Linking). doi: 10.1038/s41591-021-01318-5. URL https://www.ncbi.nlm.nih.gov/pubmed/33772244.

[23] W. F. Garcia-Beltran, E. C. Lam, K. St Denis, A. D. Nitido, Z. H. Garcia, B. M. Hauser, J. Feldman, M. N. Pavlovic, D. J. Gregory, M. C. Poznansky, A. Sigal, A. G. Schmidt, A. J. Iafrate, V. Naranbhai, and A. B. Balazs. Multiple sars-cov-2 variants escape neutralization by vaccine-induced humoral immunity. Cell, 184(9):2523, 2021. ISSN 1097-4172 (Electronic) 0092-8674 (Linking). doi: 10.1016/j.cell.2021.04.006. URL https://www.ncbi.nlm.nih.gov/pubmed/33930298.

[24] S. A. Madhi, V. Baillie, C. L. Cutland, M. Voysey, A. L. Koen, L. Fairlie, S. D. Padayachee, J. Dheda, S. L. Barnabas, Q. E. Bhorat, C. Briner, G. Kwatra, K. Ahmed, P. Aley, S. Bhikha, J. N. Bhiman, A. E. Bhorat, J. du Plessis, A. Esmail, M. Groenewald, E. Horne, S. H. Hwa, A. Jose, T. Lambe, M. Laubscher, M. Malahleha, M. Masenya, M. Masilela, S. McKenzie, K. Molapo, A. Moultrie, S. Oelofse, F. Patel, S. Pillay, S. Rhead, H. Rodel, L. Rossouw, C. Taoushanis, H. Tegally, A. Thombrayil, S. van Eck, C. K. Wibmer, N. M. Durham, E. J. Kelly, T. L. Villafana, S. Gilbert, A. J. Pollard, T. de Oliveira, P. L. Moore, A. Sigal, A. Izu, and Ngs-Sa Group Wits-VIDA COVID Group. Efficacy of the chadox1 ncov-19 covid-19 vaccine against the b.1.351 variant. N Engl J Med, 2021. ISSN 1533-4406 (Electronic) 0028-4793 (Linking). doi: 10.1056/NE-JMoa2102214. URL https://www.ncbi.nlm.nih.gov/pubmed/33725432.

[25] X. Xie, Y. Liu, J. Liu, X. Zhang, J. Zou, C. R. Fontes-Garfias, H. Xia, K. A. Swanson, M. Cutler, D. Cooper, V. D. Menachery, S. C. Weaver, P. R. Dormitzer, and P. Y. Shi. Neutralization of SARS-CoV-2 spike 69/70 deletion, E484K and N501Y variants by BNT162b2 vaccine-elicited sera. Nat Med, 2021. ISSN 1546-170X (Electronic) 1078-8956 (Linking). doi: 10.1038/s41591-021-01270-4. URL https://www.ncbi.nlm.nih.gov/pubmed/33558724.

[26] Z. Wang, F. Schmidt, Y. Weisblum, F. Muecksch, C. O. Barnes, S. Finkin, D. Schaefer-Babajew, M. Cipolla, C. Gaebler, J. A. Lieberman, T. Y. Oliveira, Z. Yang, M. E. Abernathy, K. E. Huey-Tubman, A. Hurley, M. Turroja, K. A. West, K. Gordon, K. G. Millard, V. Ramos, J. D. Silva, J. Xu, R. A. Colbert, R. Patel, J. Dizon, C. Unson-O’Brien, I. Shimeliovich, A. Gazumyan, M. Caskey, P. J. Bjorkman, R. Casellas, T. Hatziioannou, P. D. Bieniasz, and M. C. Nussenzweig. mRNA vaccine-elicited antibodies to SARS-CoV-2 and circulating variants. Nature, 2021. ISSN 1476-4687 (Electronic) 0028-0836 (Linking). doi: 10.1038/s41586-021-03324-6. URL https://www.ncbi.nlm.nih.gov/pubmed/33567448.

[27] K. Wu, A. P. Werner, M. Koch, A. Choi, E. Narayanan, G. B. E. Stewart-Jones, T. Colpitts, H. Bennett, S. Boyoglu-Barnum, W. Shi, J. I. Moliva, N. J. Sullivan, B. S. Graham, A. Carfi, K. S. Corbett, R. A. Seder, and D. K. Edwards. Serum neutralizing activity elicited by mRNA-1273 vaccine – preliminary report. N Engl J Med, 2021. ISSN 1533-4406 (Electronic) 0028-4793 (Linking). doi: 10.1056/NEJMc2102179. URL https://www.ncbi.nlm.nih.gov/pubmed/33596346.

[28] Markus Hoffmann, Prerna Arora, Rüdiger Groß, Alina Seidel, Bojan Hörnich, Alexander Hahn, Nadine Krüger, Luise Graichen, Heike Hofmann-Winkler, Amy Kempf, Martin Sebastian Winkler, Sebastian Schulz, Hans-Martin Jäck, Bernd Jahrsdörfer, Hubert Schrezenmeier, Martin Müller, Alexander Kleger, Jan Münch, and Stefan Pöhlmann. SARS-CoV-2 variants B.1.351 and B.1.1.248: Escape from therapeutic antibodies and antibodies induced by infection and vaccination. bioRxiv, 2021. doi: 10.1101/2021.02.11.430787. URL https://www.biorxiv.org/content/early/2021/02/11/2021.02.11.430787.

[29] T. Skelly Donal, C. Harding Adam, Gilbert-Jaramillo Javier, L. Knight Michael, Longet Stephanie, Anthony Brown, Adele Sandra, Adland Emily, Brown Helen, Team Medawar Laboratory, Tom Tipton, Stafford Lizzie, A. Johnson Síle, Amini Ali, Optic Clinical Group, Kit Tan Tiong, Schimanski Lisa, A. Huang Kuan-Ying, Rijal Pramila, Pitch Study Group, Cmore Phosp-C. Group, Frater John, Goulder Philip, P. Conlon Christopher, Jeffery Katie, Dold Christina, J. Pollard Andrew, R. Townsend Alain, Klenerman Paul, J. Dunachie Susanna, Barnes Eleanor, W. Carroll Miles, and S. James William. Research Square, 2021. ISSN 2693-5015. doi: 10.21203/rs.3.rs-226857/v1. URL https://doi.org/10.21203/rs.3.rs-226857/v1.

[30] P. Wang, L. Liu, S. Iketani, Y. Luo, Y. Guo, M. Wang, J. Yu, B. Zhang, P. D. Kwong, B. S. Graham, J. R. Mascola, J. Y. Chang, M. T. Yin, M. Sobieszczyk, C. A. Kyratsous, L. Shapiro, Z. Sheng, M. S. Nair, Y. Huang, and D. D. Ho. Increased resistance of SARS-CoV-2 variants B.1.351 and B.1.1.7 to antibody neutralization. bioRxiv, 2021. doi: 10.1101/2021.01.25.428137. URL https://www.ncbi.nlm.nih.gov/pubmed/33532778.

[31] T. Moyo-Gwete, M. Madzivhandila, Z. Makhado, F. Ayres, D. Mhlanga, B. Oosthuysen, B. E. Lambson, P. Kgagudi, H. Tegally, A. Iranzadeh, D. Doolabh, L. Tyers, L. R. Chinhoyi, M. Mennen, S. Skelem, G. Marais, C. K. Wibmer, J. N. Bhiman, V. Ueckermann, T. Rossouw, M. Boswell, T. de Oliveira, C. Williamson, W. A. Burgers, N. Ntusi, L. Morris, and P. L. Moore. Cross-reactive neutralizing antibody responses elicited by sars-cov-2 501y.v2 (b.1.351). N Engl J Med, 2021. ISSN 1533-4406 (Electronic) 0028-4793 (Linking). doi: 10.1056/NEJMc2104192. URL https://www.ncbi.nlm.nih.gov/pubmed/33826816.

[32] James Brett Case, Adam L Bailey, Arthur S Kim, Rita E Chen, and Michael S Diamond. Growth, detection, quantification, and inactivation of SARS-CoV-2. Virology, 2020.

[33] Alex Sigal, Tamar Danon, Ariel Cohen, Ron Milo, Naama Geva-Zatorsky, Gila Lustig, Yuvalal Liron, Uri Alon, and Natalie Perzov. Generation of a fluorescently labeled endogenous protein library in living human cells. Nature protocols, 2(6):1515–1527, 2007.

[34] Y. M. Bar-On, A. Flamholz, R. Phillips, and R. Milo. Sars-cov-2 (covid-19) by the numbers. Elife, 9, 2020. ISSN 2050-084X (Electronic) 2050-084X (Linking). doi: 10.7554/eLife.57309. URL https://www.ncbi.nlm.nih.gov/pubmed/32228860.

[35] Y. Liu, J. Liu, H. Xia, X. Zhang, C. R. Fontes-Garfias, K. A. Swanson, H. Cai, R. Sarkar, W. Chen, M. Cutler, D. Cooper, S. C. Weaver, A. Muik, U. Sahin, K. U. Jansen, X. Xie, P. R. Dormitzer, and P. Y. Shi. Neutralizing activity of BNT162b2-elicited serum – preliminary report. N Engl J Med, 2021. ISSN 1533-4406 (Electronic) 0028-4793 (Linking). doi: 10.1056/NEJMc2102017. URL https://www.ncbi.nlm.nih.gov/pubmed/33596352.

[36] D. F. Robbiani, C. Gaebler, F. Muecksch, J. C. C. Lorenzi, Z. Wang, A. Cho, M. Agudelo, C. O. Barnes, A. Gazumyan, S. Finkin, T. Hagglof, T. Y. Oliveira, C. Viant, A. Hurley, H. H. Hoffmann, K. G. Millard, R. G. Kost, M. Cipolla, K. Gordon, F. Bianchini, S. T. Chen, V. Ramos, R. Patel, J. Dizon, I. Shimeliovich, P. Mendoza, H. Hartweger, L. Nogueira, M. Pack, J. Horowitz, F. Schmidt, Y. Weisblum, E. Michailidis, A. W. Ashbrook, E. Waltari, J. E. Pak, K. E. Huey-Tubman, N. Koranda, P. R. Hoffman, Jr. West, A. P., C. M. Rice, T. Hatziioannou, P. J. Bjorkman, P. D. Bieniasz, M. Caskey, and M. C. Nussenzweig. Convergent antibody responses to SARS-CoV-2 in convalescent individuals. Nature, 584(7821):437–442, 2020. ISSN 1476-4687 (Electronic) 0028-0836 (Linking). doi: 10.1038/s41586-020-2456-9. URL https://www.ncbi.nlm.nih.gov/pubmed/32555388.

[37] A. Tarke, J. Sidney, C. K. Kidd, J. M. Dan, S. I. Ramirez, E. D. Yu, J. Mateus, R. da Silva Antunes, E. Moore, P. Rubiro, N. Methot, E. Phillips, S. Mallal, A. Frazier, S. A. Rawlings, J. A. Greenbaum, B. Peters, D. M. Smith, S. Crotty, D. Weiskopf, A. Grifoni, and A. Sette. Comprehensive analysis of t cell immunodominance and immunoprevalence of SARS-CoV-2 epitopes in Covid-19 cases. Cell Rep Med, 2(2):100204, 2021. ISSN 2666-3791 (Electronic) 2666-3791 (Linking). doi: 10.1016/j.xcrm.2021.100204. URL https://www.ncbi.nlm.nih.gov/pubmed/33521695.

